# Auditory attention reduced ear-canal noise in humans, but not through medial olivocochlear efferent inhibition: Implications for measuring otoacoustic emissions during behavioral task performance

**DOI:** 10.1101/354902

**Authors:** Nikolas A. Francis, Wei Zhao, John J. Guinan

## Abstract

Otoacoustic emissions (OAEs) are often measured to non-invasively determine activation of medial olivocochlear (MOC) efferents in humans. Usually these experiments assume that ear-canal noise remains constant. However, changes in ear-canal noise have been reported in some behavioral experiments. We studied the variability of ear-canal noise in eight subjects who performed a two-interval-forced-choice (2IFC) sound-level-discrimination task on monaural tone pips in masking noise. Ear-canal noise was recorded directly from the unstimulated ear opposite the task ear. Recordings were also done with similar sounds presented, but no task done. In task trials, ear-canal noise was reduced at the time the subject did the discrimination, relative to the noise level earlier in the trial. In two subjects, there was a decrease in ear-canal noise, primarily at 1-2 kHz, with a time course similar to that expected from inhibition by MOC activity elicited by the task-ear masker noise. These were the only subjects with spontaneous OAEs (SOAEs). We hypothesize that the SOAEs were inhibited by MOC activity elicited by the task-ear masker. Based on the standard rationale in OAE experiments that large bursts of noise are artifacts due to subject movement, noise bursts above a sound-level criterion were removed. As the criterion was lowered and more high-and moderate-level noise bursts were removed, the reduction in noise level from the beginning of the trial to the time of the 2IFC discrimination became less. This pattern is opposite that expected from MOC inhibition (which is greater on lower-level sounds), but can be explained by the hypothesis that subjects move less and create fewer bursts of noise when they concentrate on doing the task. In contrast, for the six subjects with no SOAEs, in no-task trials the noise level was little changed throughout the trial. Our results show that measurements of MOC effects on OAEs must measure and account for changes in ear-canal noise, especially in behavioral experiments. The results also provide a novel way of showing the time course of the buildup of attention in ear-canal noise during a 2IFC task.

## 1 Introduction

Medial olivocochlear (MOC) efferent activity has long been hypothesized to facilitate hearing in noise (Nieder and Nieder, 1970; Michel and Collet, 1993; Guinan, 1996). Many papers have attempted to determine how MOC efferent activity affects hearing by measuring changes in otoacoustic emissions (OAEs) as subjects performed an auditory task that was expected to elicit efferent activity (e.g. Puel et al., 1988; Meric and Collet, 1994; de Boer and Thornton, 2007; Harkrider and Bowers). MOC activity reduces the gain of cochlear amplification and thereby reduces OAEs, so OAE reductions provide information about efferent activation and its effects in the cochlea. A key assumption in measuring OAEs during behavioral task performance has been that there is no change in the background level of the random noise in the ear canal, so that any measured changes in OAEs can be attributed to changes produced by MOC efferents.

In contrast to the assumption that ear-canal noise is not changed during a behavioral task, several studies have reported such changes (de Boer and Thornton, 2007; Walsh et al., 2014a; 2014b, 2015). Walsh et al. (2014; 2015) reported that ear-canal random noise was reduced by selective attention activating MOC efferents. In the Walsh et al. experiments, ear-canal noise was indirectly measured during a 30 ms silent period by a double-evoked technique that yielded a measure termed a “nonlinear stimulus frequency otoacoustic emission” or “nSFOAE” (Walsh et al., 2010). During both auditory and visual tasks there was a reduction in ear-canal noise (i.e. a reduction in the nSFOAE) relative to when the subject was presented the same stimuli but did not do a task (Walsh et al., 2014a; 2014b, 2015). For an auditory task, the reduction was similar in both the attended ear and the opposite ear. Walsh et al. hypothesized that cochlear-amplified random vibrations within the cochlea created backward traveling waves that produced acoustic noise in the ear canal, and activation of MOC efferents reduced cochlear amplification and therefore reduced the random noise within the ear canal.

We have done experiments that allow us to measure changes in ear-canal acoustic noise during a behavioral task. Our subjects did a two-interval-forced-choice (2IFC) level discrimination task on monaural tone bursts in noise. During these tests we measured changes in click-evoked otoacoustic emissions (CEOAEs) in the task ear, with the goal of assessing changes in MOC activation during the behavioral task. Most relevant here is that we also measured the sound pressure in the ear where no sound was presented, opposite to the task ear. These opposite-ear recordings provide an opportunity to directly determine whether there was a reduction in ear-canal background noise sound pressure during the behavioral task, and to measure its time course relative to the time when sounds were presented and the subject made the 2IFC judgment.

## 2 Methods

### 2.1 Subjects

Eight subjects (aged 18-21 years; 2 male) participated in the experiments reported here. All subjects had normal pure-tone audiograms (<15 dB HL at octave frequencies 0.5 to 8 kHz). Sounds were presented and recorded using Etymotic Research ER10c acoustic assemblies, sampled at 25 kHz. The acoustic outputs were monitored and calibrated frequently throughout the experiments. This study was performed in accordance with MEEI, MIT and NIH guidelines for human studies. Informed consent was obtained from all subjects.

### 2.2 Experimental methods

The experiments were designed to detect changes in CEOAE amplitudes brought about by efferent activity, i.e. changes in CEOAEs from the beginning of a 2IFC trial to just after the stimuli to be discriminated in the trial (the masking noise made it too difficult to measure CEOAEs during the noise). We did both “active” runs in which the subject did the 2IFC task, and “passive” runs in which the subject heard the same sounds but made no judgment. Since learning to do the 2IFC task might cause a subject to continue to attend to the task sounds during passive trials, passive trials were done first, before the subjects were told about their future task. Passive and active conditions were typically done in separate sessions, where a “session” is defined as the time that a subject continuously had the acoustic-assembly foam plugs in their ear canals. Removing and replacing the acoustic assembly was considered a new session, whether it was a few minutes later or days later. Since acoustic parameters such as the depth of insertion might change across sessions, direct comparisons of the amplitudes of the ear-canal acoustic noise in active versus passive listening were not done because such comparisons may not be accurate. However, the stimuli and their timing were the same in passive and active trials so we can compare the time courses of ear-canal sound in passive and active trials.

Sound stimuli were presented only in the task ear, which was the ear that had the most robust CEOAEs in our initial tests. The subject’s task was to detect which of two short tone bursts was larger in amplitude. Both tone bursts were embedded in 50 dB SPL broad-band noise. The baseline level of the tone bursts (the pedestal level) was varied between sessions and set to no-pedestal, 40, 50, 60, 70 or 80 dB SPL. The two tone bursts were stepped in level about the pedestal level (one up, one down) by the same number of dB (or just up for no-pedestal, i.e., a tone was presented only in one interval). The tone burst with the higher level was chosen randomly on each trial. For each subject and pedestal level, the step size was chosen to achieve a correct response rate of 84%. In passive trials the step size was zero.

Data were collected in batches of 25 trials, with the same pedestal level throughout the batch. On each trial, sound was presented only in the task ear in a continuous series of 400 ms epochs with 50 dB pSPL, 80 μs rarefaction clicks at 25 ms intervals presented throughout each epoch (16 clicks per epoch). Each trial began with 1 to 10 epochs (number randomly selected on each trial) containing only clicks (see Fig. 1). This was followed by 3 epochs that had the clicks plus 50 dB SPL, broadband (0.1-10 kHz) frozen noise (the same in each epoch). The last two epochs with clicks and noise also had a tone burst (15 ms plateau, 5 ms raised-cosine rise and fall times) that ended 45 ms before the end of the epoch. After the tone-in-noise epochs there was an additional 400 ms epoch in which there were only repeated clicks (the same as in initial epochs 1-10) (Fig. 1). Overall, the number of 400 ms epochs in each trial varied from 5 to 14, depending on the number of initial epochs. At the end of each trial the subject indicated whether the first or second tone burst was higher in level by pushing one of two buttons on a device on which their hand rested (usually this done was during the last 400 ms epoch). To push the proper button, a subject only had to move one finger and did not have to move their arm. We did not have subjects type on a keypad or touch a screen so as to minimize subject motion. The next trial in the batch of 25 trials began 1 second after the button press or end of the last epoch, whichever came later.

**Figure 1.**
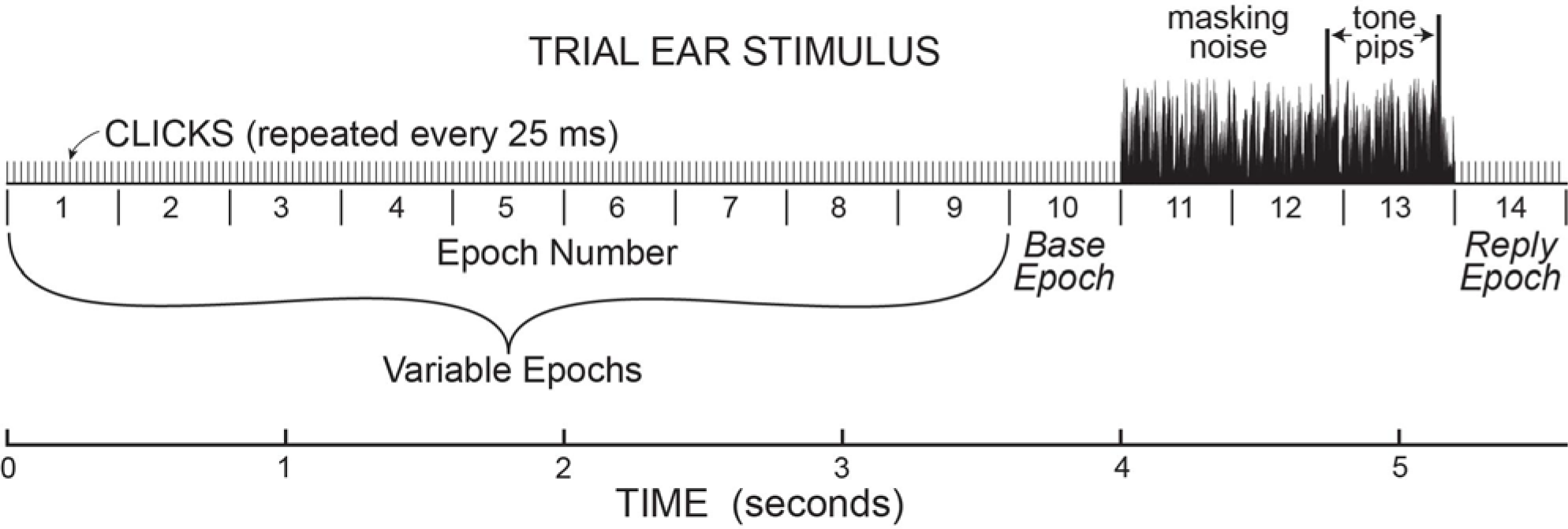
Stimulus paradigm for the sounds presented in the task ear in one trial. This paradigm established the timing for the analysis of ear-canal noise in the opposite ear, where no sound was presented. In the task ear, stimuli were presented in 400 ms “epochs” with 50 dB pSPL clicks presented every 25 ms throughout. There were 1-10 (randomly chosen) epochs before three epochs with 50 dB SPL masking noise, followed by one epoch at the end. The epochs from “base” to “reply” were present on every trial. The last two epochs with noise (epochs 12&13) also contained tone pips, which were the same amplitude in the no-task trials, and different amplitude in task trials. At the end of each trial during the task, subject had to push a button to indicate which tone pip was louder.

Spontaneous otoacoustic emissions (SOAEs) were measured once on each subject by recording the ear canal sound in both ears simultaneously with no stimulus presented and the subjects instructed to sit very still for this short measurement. On each ear, eight data buffers were obtained, each sampled every 40 μs and 2.62 seconds long. Each buffer was individually fast-Fourier transformed and the resulting amplitudes (phases set to zero) were averaged. Two subjects (323, 326) had SOAEs, as judged by their having spectral lines that were >10 dB above the smoothed SOAE spectra.

### 2.3 Data analysis

Throughout each trial, sound was recorded continuously in both ears and stored for later processing. The data for the present paper are from the ear opposite the task ear, except that the test for middle-ear-muscle (MEM) activation used the amplitudes of the clicks in the task ear. Before processing, the opposite-ear data were filtered from 0.5-5 kHz by a zero-phase-change FIR digital filter. The opposite-ear recordings were divided into 25 ms time spans—hereafter referred to as “spans”— corresponding to the times demarcated by the clicks in Figure 1. We measured the root-mean-square (RMS) value of the sound in every time span. We visualized the amplitude distribution of the RMS’s from the spans in a batch of 25 trials—hereafter referred to as a “batch”—by binning the RMS values into 300-bin histograms with the 100^th^ bin equal to the median value of the RMS distribution and bin widths of 1% of the median value (Fig. 2). RMS values greater than three times the median value were used later, but were omitted from the histogram. For most sessions, these RMS histograms had narrow peaks and tails with higher RMS values (e.g. Fig. 2). For subsequent data analysis, a given span was not used if its RMS value was above a rejection criterion RMS value that was a parameter varied in our study. To find a criterion value, we first smoothed the histogram and then determined an “upper-edge RMS” value, where the histogram fell to 50% of the peak. The difference between the upper-edge-RMS and the peak RMS is termed the “Edge Width”. The Edge Width, multiplied by a user-chosen constant (the “Edge Multiplier), and added to the peak RMS value, defined the rejection criterion RMS value.

**Figure 2.**
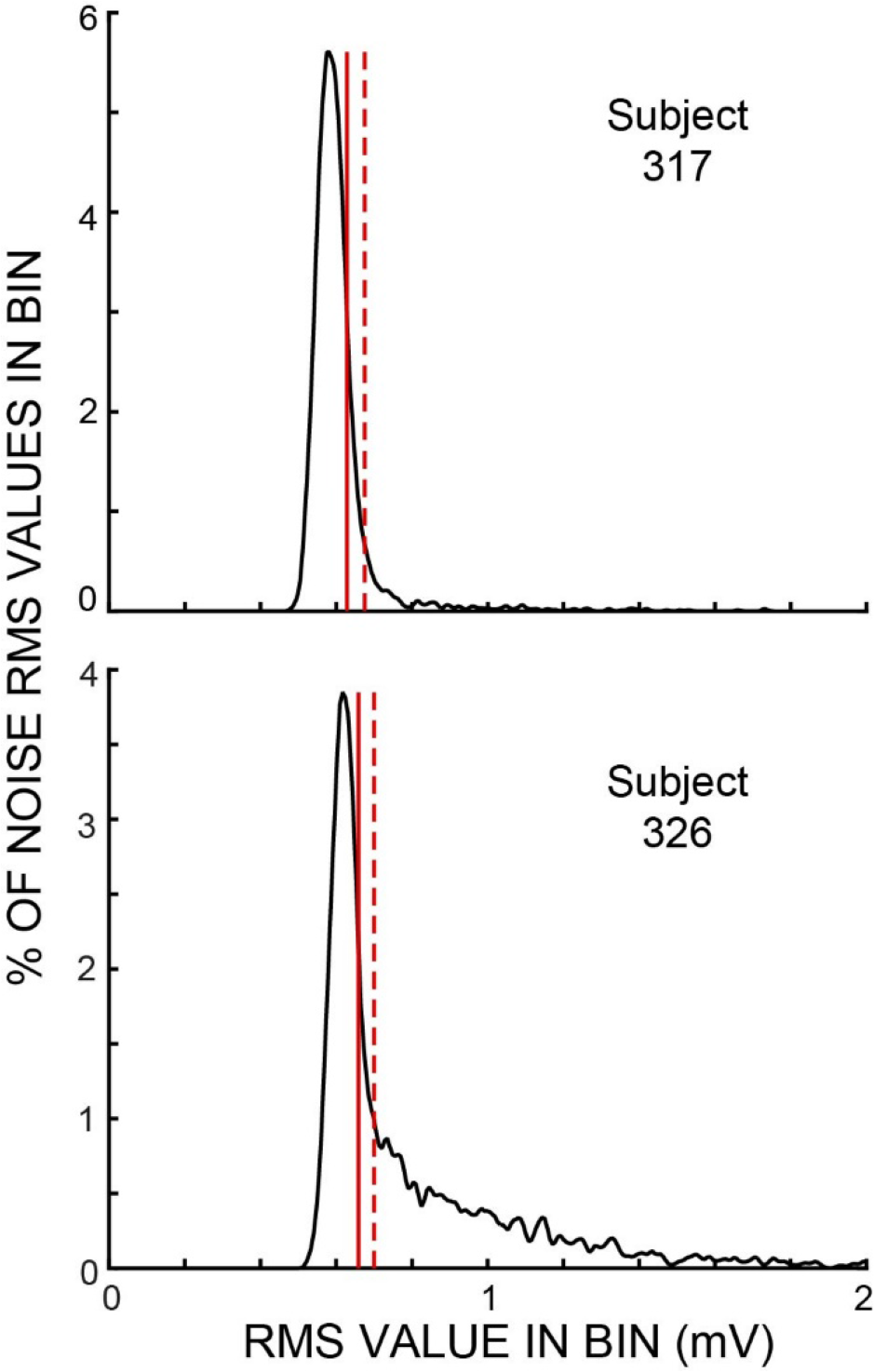
Histograms of span RMS values from one batch of trials for each of two subjects for the epochs without masker noise. These subjects were chosen to show different amounts of noise at sound levels above the peak region. Vertical solid lines show the “upper edge” at which the histograms fell to 50% of the peak. Vertical dashed lines show the sound-level cut off for a noise rejection criterion value (i.e., an edge multiplier - see Methods) of 2. Both examples show data from active trials. In these two batches, the peaks were at 17.2 dB SPL (subject 317) and 15.4 dB SPL (subject 326).

The opposite-ear sound recordings were contaminated to varying degrees by crosstalk from the task-ear masker noise. This crosstalk was assessed from the difference between two ways of combining pairs of span waveforms from different trials of a batch: (1) reversing the polarity of one waveform of the pair and then averaging, or (2) averaging the waveforms without reversing either one (Fig. 3). Since the frozen-noise masker was the same on every trial, reversing the polarity of one waveform before averaging cancels the crosstalk contribution in the average. In contrast, if the ear-canal sound is random noise, reversing the sign of a waveform before averaging makes no difference. The difference in these two measures (each averaged over the time when the masker noise was on: epochs 11-13) and converted to dB SPL, yielded crosstalk levels averaging −22 dB SPL (range −31 to −10 dB SPL). We compensated for the square-root-of-2 adjustment appropriate for averaging noise but not appropriate for averaging the crosstalk signal. The task-ear masker noise was 50 dB SPL so the crosstalk attenuation averaged 72 dB. In a few sessions, the crosstalk and/or other aspects of the recordings were highly abnormal (differed by more than a factor of two from the other values on that subject - perhaps the acoustic assemblies were not properly seated); these data were excluded from our analysis.

**Figure 3.**
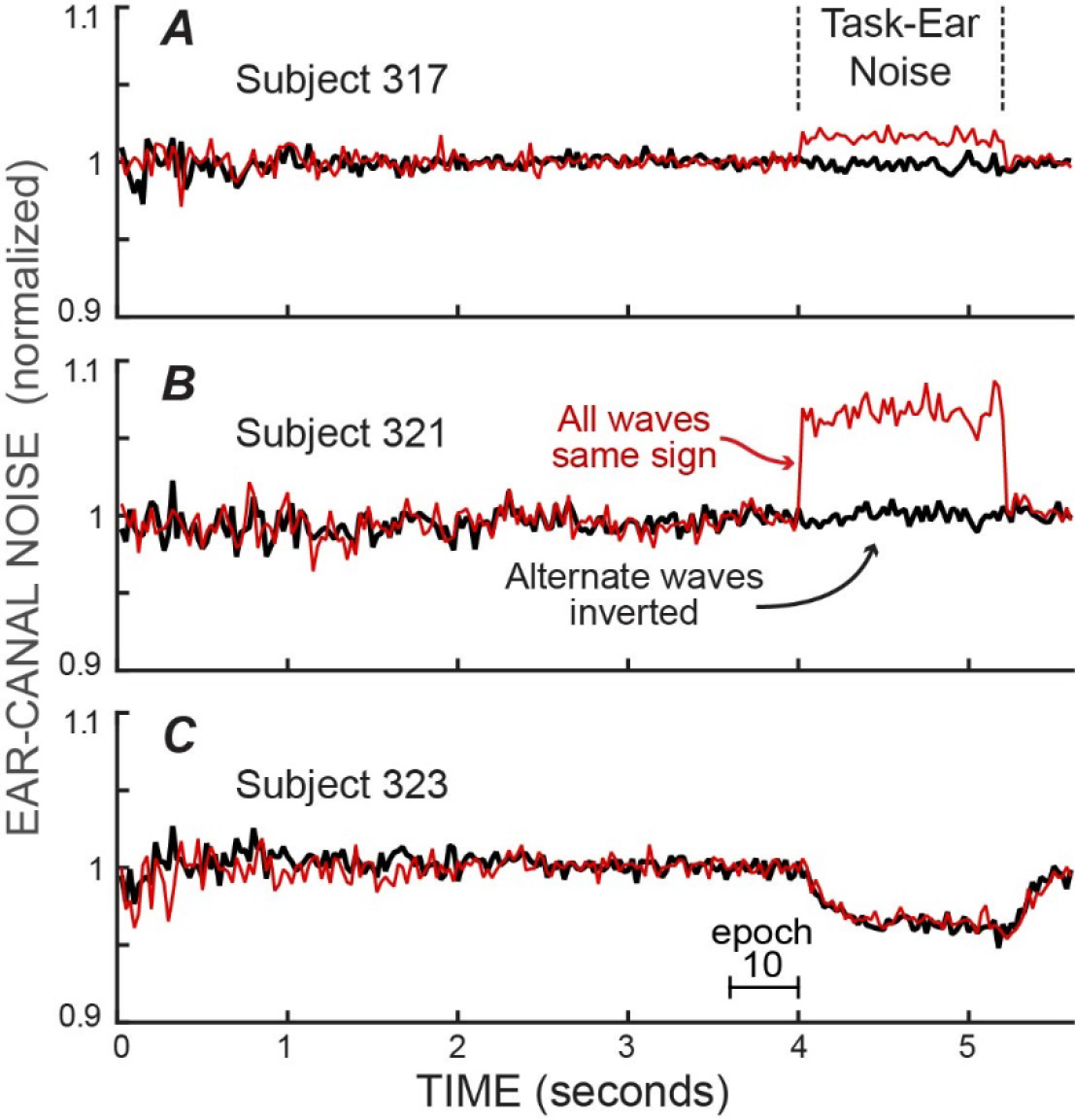
The RMS values of the ear-canal noise in successive 20 ms spans, normalized by the value in epoch 10, showing the effect of crosstalk from the task-ear frozen-noise masker. The effect of the crosstalk can be seen by comparing: (1) the averaged pairs of recordings with both waves the same sign, which averages the crosstalk (thin, light traces), versus (2) the averaged pairs after inverting one of the pair before averaging, which cancels the crosstalk (thick, dark traces). The differences during the task-ear noise show that there was almost no crosstalk in C, little crosstalk in A, and crosstalk that increased the ear-canal sound by about 6% in B. In addition, the traces with the crosstalk cancelled (thick, dark traces) allow the ear-canal noise during the presence of the task-ear masker to be compared with the ear-canal noise before and after the masker. No detectable decrease in ear-canal noise during the masker is seen in A and B, while the largest decrease of any subject (~4%) is seen in C. The decrease in C has a time course similar to that expected from masker-evoked medial olivocochlear (MOC) efferent inhibition.

To avoid masker-noise crosstalk from affecting the noise rejections, we used a two-step procedure to exclude noisy spans. The procedure described below was applied separately for each of the spans that occurred at a given time in a batch, whether or not the span was from the time when the masker noise was present. First, individual spans were excluded if their RMS level was above a rejection criterion that was twice as far from the peak as the regular criterion (i.e. we used two times the value of the edge multiplier). This removed spans with particularly large-amplitude noises that would be rejected no matter how low the noise was in any span they would be paired with. Spans that passed this first criterion were paired by summing their waveforms point-by-point with one of the pair reversed in polarity (to cancel the crosstalk) and from the summed waveform we calculated the reverse-pair-RMS value. Next, data from such a pair were excluded if the reverse-pair-RMS was above the rejection criterion multiplied by the square-root of two to compensate for adding orthogonal noise waveforms. The reverse-pair-RMS’s of all the passed pairs in a batch were summed, and the sum was divided by the number of spans that passed the rejection criteria. This yielded a single average RMS value for the noise of a span in a given batch. This was done separately for successive spans across the 14 epochs, yielding a time-course of RMS values across a trial.

The RMS values for each span in a batch were expressed in two ways: (1) RMS values were converted to a linear version of dB SPL (“linear SPL”) using the appropriate acoustic calibration. These averages, converted to dB, were used when plotting the amplitudes in dB SPL. (2) RMS values were normalized by dividing each span by the average RMS value of the spans in epoch 10 of the batch. For each method, data at each successive span were combined across batches by averaging the RMS values. Batches were identified as “active” or “passive” and were averaged separately. In some subjects, crosstalk sound from the highest pedestal levels was not canceled by averaging alternated-sign waveform pairs because the tone bursts randomly varied in amplitude so that adjacent waveforms did not always have the same amplitude tone burst and therefore did not cancel. Data from these pedestal levels were excluded from plots (31%, on average, including all of the 80 dB pedestals); otherwise differences in pedestal level were ignored because we found no systematic differences in ear-canal noise levels from batches with different tone burst pedestal levels.

The resulting span RMS values, and the fraction of spans rejected, were plotted across time in successive 25-ms time spans. Although individual trials had different numbers of initial epochs, we used a timing scheme for displaying the data in which the first time span plotted in a time course was chosen as if every trial had all 10 of the initial 400 ms stimulus epochs. Spans in the first epoch had actual sound-recording data only in ~10% of trials, since we randomized of the number of initial epochs (1-10) for each trial. Spans in epochs 2 to 10 each had data in successively 10% more trials. Spans in epochs 10 to 14 had sound-recording data in all trials. Overall there were 224 spans (14 epochs multiplied by 16 spans per epoch) with the last one ending at the end of the final epoch.

The middle-ear-muscle (MEM) reflex is bilateral, so the masker noise in the task-ear may have elicited MEM contractions that could affect the noise in the opposite ear. We tested for MEM contractions on each trial by comparing the click amplitudes in the task ear before and after the masking-noise epochs. MEM contractions stiffen the ossicular chain which typically increases the ear-canal sound pressure produced by a constant sound source. In each trial, we averaged click amplitudes throughout epoch 10 and also averaged 12 clicks of epoch 14 starting with the second click (in epoch 14, the first click was contaminated by effects of the masker noise and later clicks were not used to avoid times after MEM contractions would have decayed). If the increase in click amplitude exceeded 0.2 dB, data from that trial were not used. With this criterion, data from ~0.5 to 4% of trials across subjects were excluded. However, because the rejected trials were not systematically from certain subjects or pedestal levels, we think these rejected trials were not due to actual MEM contractions.

The spectra of the ear-canal noise were obtained by a filter-bank method similar to that used by Francis and Guinan (2010). We used zero-phase-change FIR digital filters. Individual filters were 500 Hz wide, with center frequencies 250 Hz apart (they overlapped), and extending from 500 to 4000 Hz. Span waveform pairs were combined with one of the pair reversed, so as to cancel any crosstalk. They were accepted or rejected by their RMS values as described above, and then each accepted pair was filtered to obtain its spectra. For each subject, span spectra were combined by averaging in 6 groups: for epochs 1-9, all spectra were combined in a single group, and for epochs 10 to 14, all of the spectra from each epoch were combined into separate epoch averages. In all cases, spectra from active and passive trials were combined separately.

### 2.4 Statistical analysis

To determine if changes in ear-canal sound recordings were statistically significant, we used a bootstrap test (an ANOVA could not be used because the data were not normally distributed, see Fig. 2). Bootstrap tests were applied separately on each subject and each activity group (active or passive) using averages of the span data in epochs 10 to 14 (each epoch averaged separately). Separate tests were done for the normalized noise and for the fraction rejected. For each set of data, all of the batches included in the original average for that group (N batches averaging 37.8, range 20 to 59 across subjects), formed the set of input batches for the bootstrap. From the N batches of a set, new sets of N batches were formed by randomly selecting a batch from the original set (but not removing it from the original set so it could be selected again), and doing this N times. For each new set of N batches, new epoch averages for epochs 10 to 14 were calculated in the same way as for the original calculation. After averaging, the data from epochs 11 to 14 were each normalized relative to the data in epoch 10 by dividing by the value in epoch 10 for noise amplitudes or by subtracting for the fraction of spans rejected. New sets of N batches were obtained 100,000 times, which yielded 100,000 new averages for each epoch. With the hypothesis that the average noise level in each of epochs 11 to 14 was smaller than in epoch 10, the fraction of times that a normalized epoch average was *higher* than unity is the probability that the hypothesis was false, i.e. this is the significance level (the p value) for the hypothesis that the average value in a given epoch from 11 to 14 was less than the average value in epoch 10.

To compare whether the reduction in the ear-canal noise from epoch 10 to epochs 11-14 was more in the active trials than in the passive trials of a subject, new pseudo-average values of the changes from epoch 10 to epochs 11-14 were calculated separately for the active and passive trials as described above. We calculated the noise reduction as: (epoch 10 - epoch X). From these new pseudo-averages, for each epoch we calculated the additional reduction of the ear-canal noise in the active trials compared to the passive trials (i.e. the active value minus the passive value) and if this value was less than zero, the comparison was scored as false. This was done 100,000 times and the fraction false was taken as the probability that the hypothesis was false. This is the p value for the hypothesis that the reduction of ear-canal noise from epoch 10 to epochs 11-14 was more in the active trials than in the passive trials.

## 3 Results

### 3.1 No noise rejections

Ear-canal noise levels, expressed as dB SPL values in successive 25 ms time spans (Fig. 4A, B), were measured when the subjects were doing the 2IFC task (active trials) and when subjects sat quietly without doing the task (passive trials). The overall noise levels varied across subjects and overlapped considerably. To make the trends easier to see, each set of data was normalized (using SPLs as linear numbers) relative to their average value in the base epoch (epoch 10) and is replotted in Figure 4C, D. In both active and passive trials the noise levels bounced around baselines that remained relatively constant until the beginning of the epochs with masking noise, i.e. starting at 4 seconds in Figure 4. After the noise onset, the active and passive trials showed different behavior. In the active trials, the noise level *decreased* near the time when the masking noise started (Fig. 4C). In contrast, in the passive trials there was no clear trend (Fig. 4D). These data show there is a big difference in the active versus the passive trials that first occurred when the subject had to attend to doing the task. It shows that the overall noise level was strongly influenced by whether the subject was doing the task, or not. This difference is present in the data without any data processing. However, it is well known that subject movement can produce noise that is picked up by an ear-canal microphone, and that subjects never sit completely still. Thus, a hypothesis that may account for these data is that the subject sat more still when paying attention to doing the task.

**Figure 4.**
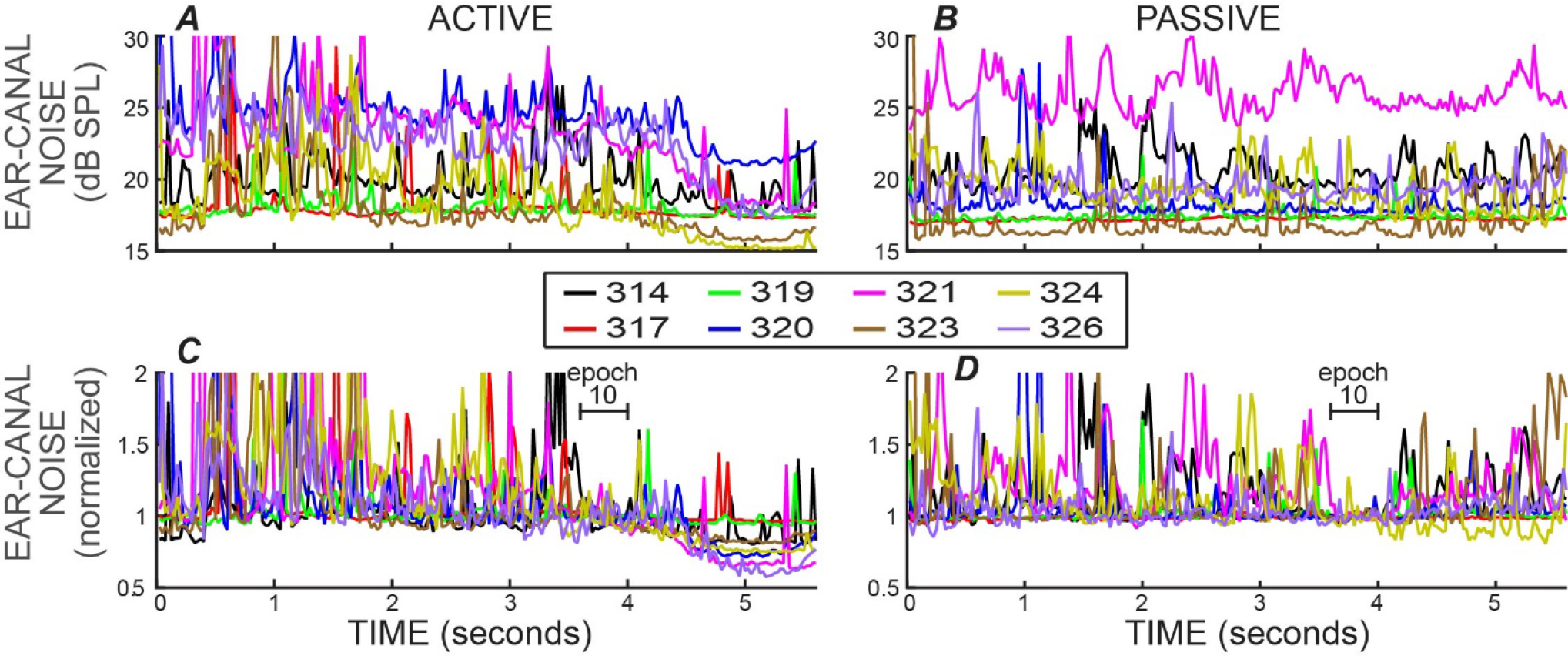
Ear-canal noise in successive 20 ms time spans for eight subjects (key in box) when the subjects were doing the task (ACTIVE) and when they were not (PASSIVE). ***A, B***: Span RMS values in dB SPL. ***C, D***: The data of A & B normalized by dividing each subject’s data by its average value in epoch 10 (from 3.6 to 4 s).

### 3.2 Strict Noise Rejections

In almost all experiments in which OAEs are measured, an artifact rejection system is used in which the experimenter chooses a sound level criterion and portions of the recording above this criterion are removed from consideration. We used an artifact rejection system with the criteria varied by setting different “edge-multiplier” values (see Methods). For an edge-multiplier of 2, figure 5 shows example plots versus time of both the ear-canal noise and the fraction of spans rejected, for both active and passive trials. An edge-multiplier of 2 provides a strict cut off that removes all spans with RMS values above the peak region in histograms of RMS values (see Fig. 2).

**Figure 5.**
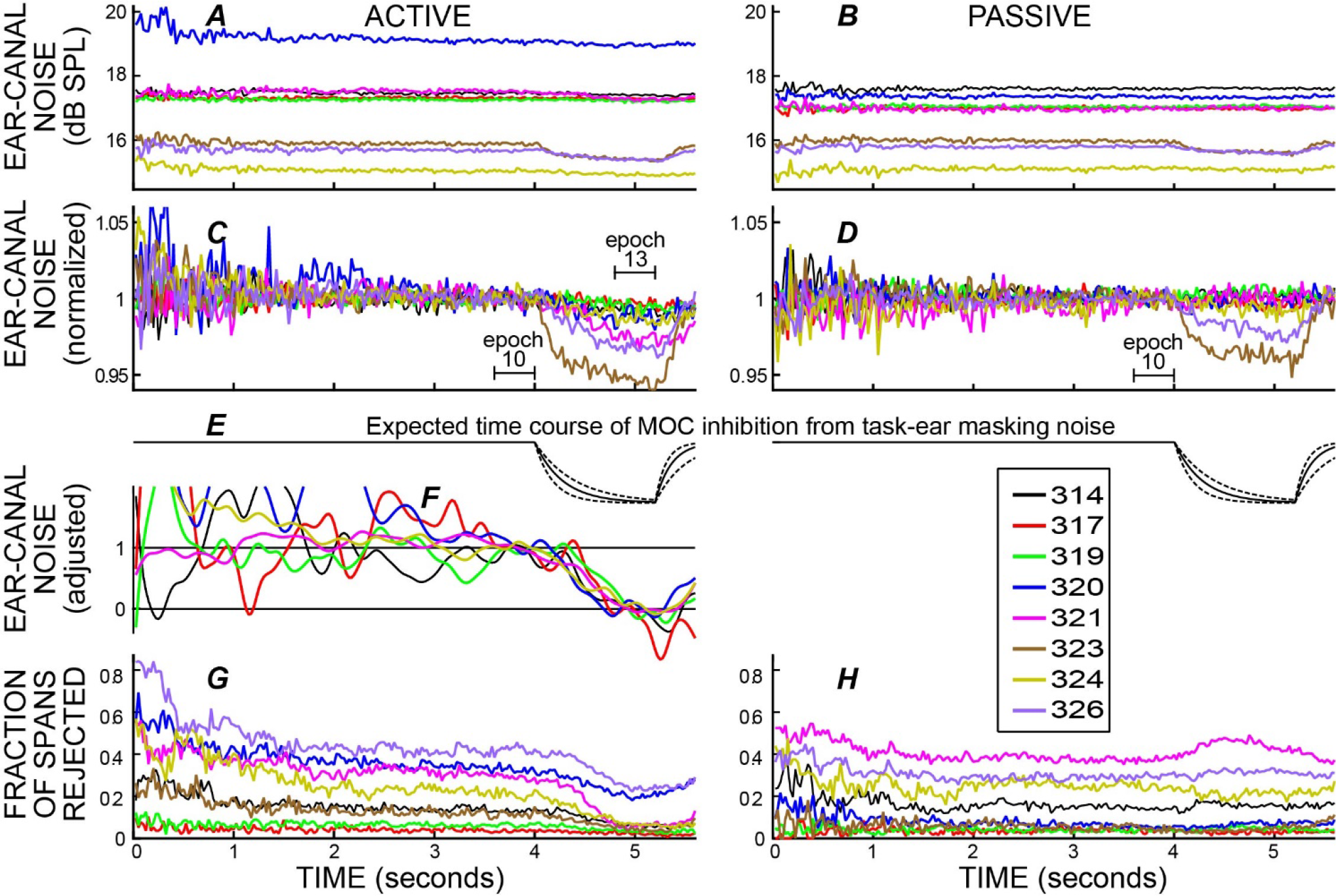
Ear-canal noise in 20 ms time spans and the fraction of spans that were rejected versus time, when subjects were doing the task (ACTIVE) and when they were not (PASSIVE) for a strict-criterion edge multiplier of 2. Data for eight subjects (key in box). ***A, B***: Span RMS values in dB SPL. ***C, D***: The data of A & B normalized by dividing each subject’s data by its average value in epoch 10. ***E***: The calculated time course of medial-olivocochlear (MOC) inhibition produced by the task-ear noise masker, based on the data of Backus and Guinan (2006). Solid lines are for the average time constants and dashed lines are for the fastest and slowest time constants. ***F***: The active-trial data from the six subjects in panels D and E who had the least change in normalized values from epoch 10 to epoch 13, with the magnification and offset adjusted so that their average in epoch 13 is zero while keeping the average in epoch 10 equal to 1. This shows the time courses of the changes independent of the amplitudes of the changes. ***G, H***: The fraction of spans rejected.

After the rejection of spans with high noise levels by applying an edge-multiplier of 2, each subject’s average noise level was relatively constant during the time before the masker noise began (Fig. 5A, B). The different SPL values for the ear-canal noise of different subjects are presumably due, at least in part, to differences in ear-canal volumes and the depths of insertion of the probes. In both active and passive trials (Fig. 5A & B), two subjects (323 and 326) had visible reductions in the overall dB SPL level of the ear-canal noise when the task-ear masker was on. These reduction are more easily seen in Figure 5C and D, which show the same data normalized to its value in epoch 10. The time courses of the decreases in ear-canal noise in these two subjects (323, 326) are similar to the time courses expected from MOC inhibition elicited by the task-ear masking noise (Fig. 5E). These two subjects were the only ones with SOAEs. A hypothesis that fits these data is that in these two subjects, the ear-canal “noise” was partly due to SOAEs that were inhibited by MOC activity elicited by the task-ear masker.

In the *passive* trials, after applying an edge multiplier of 2 to remove bursts of noise, the changes in ear-canal noise from epoch 10 to epoch 13 were all small, but some were statistically significant. The largest changes were in subjects 323 and 326 who had decreases of 3.9% and 2.4%, respectively, that were highly statistically significant (p<0.0001). In three other subjects, there were statistically significant decreases of 0.3%, 0.3% and 0.8% (p=0.016 for the least significant of these). The very small decreases in these three subjects may be due to MOC inhibition of un-noticed SOAEs or other ear-canal noise, but their time courses are too poorly defined to help substantiate this. In one subject, there was a decrease of 0.02% that was not statistically significant (p=0.47). In the two remaining subjects there were small *increases*: one increase was 0.19% but not significant (p=0.14), the other (subject 319) was an increase of 0.45% and was statistically-significant (p=0.0002).

In the *active* trials, after applying an edge-multiplier of 2 to remove bursts of noise, all of the subjects had decreases in ear-canal noise from epoch 10 to epoch 13 that were statistically-significant (p=0.00016 for the least significant). For subjects 323 and 326 the decreases were 5.4% and 3.2%, respectively, and for the six other subjects the decreases averaged 1.9% (range 0.27% to 2.6%). The largest decreases (in subjects 323 and 326) had time courses consistent with most, or all, of the decrease being from MOC inhibition elicited by the masker noise. The time courses of the decreases in the other six subjects are difficult to see in Figure 5C. To make these time courses more visible, we adjusted the magnification and offset of each so that their average in epoch 13 was zero while the average in epoch 10 was kept equal to 1. The result (Figure 5F) shows the degree to which the time courses of the reductions in ear-canal noise were similar across these six subjects. The time course of these reductions appears to have a slightly slower onset than the larger reductions seen in subjects 323 and 326 (Fig. 5C, D vs. F). However, the waveforms are somewhat noisy and the differences between them are not particularly clear.

We compared the decrease in ear-canal noise from epoch 10 to epoch 13 in active versus passive trials for an edge multiplier of 2. In 7 of 8 subjects the percentage decrease in ear-canal noise was more for the active trials than for the passive trials. The active change minus the passive change averaged 1.04%, range −0.03% to +2.5%). The greater decreases in the active trials were statistically significant in 6 of the subjects (largest p=0.005) and the one increase was not significant (p=0.56),

In addition to measuring the changes in ear-canal noise, we also measured the fraction of spans that were rejected. For an edge-multiplier of 2, the fraction of spans rejected are shown in Figure 5G, H. Near the end of the trials, when the subject had to do the 2IFC task, there was a clear difference in the fraction of spans rejected in active versus passive trials. In active trials the fraction rejected went down shortly after the masker noise started, whereas in passive trials the fraction rejected was little changed or went up (Fig. 5G, H). For active trials, all subjects had a decrease in the fraction rejected from epoch 10 to epoch 13 (average decrease = 0.107 range 0.014 to 0.23). Five of these were statistically significant (highest p=0.045) and three were not. In contrast, none of the passive trials had a statistically significant change (at the 0.05 level) in the fraction rejected in either direction over these same intervals.

Both the fractions rejected and ear-canal noise levels show the pattern over time of the bursts of noise that were present in the original data. The data of Figure 5G show that subjects reduced their production of large bursts of noise when doing the task. A hypothesis that fits these data is that large bursts of ear-canal noise are due to subject movements. With this hypothesis, the time courses of the decreases in the large-amplitude noises in Figures 4 and 5 shows the time courses over which subjects decreased movements as they directed their attention to doing the 2IFC task. In contrast, the large amplitude noises were little changed in the passive trials.

### 3.3 Varying noise rejections

The data of Figure 5 were for a strict noise-rejection criterion: an edge-multiplier of 2. For edge multipliers from 2 to 100, the reductions in ear-canal noise and the fraction of spans rejected for the active trials of all subjects are shown in Figure 6. Higher edge-multipliers reject fewer spans, but the pattern across time of the fraction of spans rejected changed little as the edge-multiplier was changed. When the criterion removed only very highest level sounds (edge multiplier of 100), or when the criterion removed all of the noise levels above the main peak in the span RMS histograms (edge multiplier of 2), the fraction rejected was lowest at the time when the subject had to make the 2IFC judgment (Fig. 6). Further, for each subject, the time courses of the reductions in ear-canal noise were very similar to the time courses of the fractions rejected, presumably because both were due to the same underlying cause.

**Figure 6.**
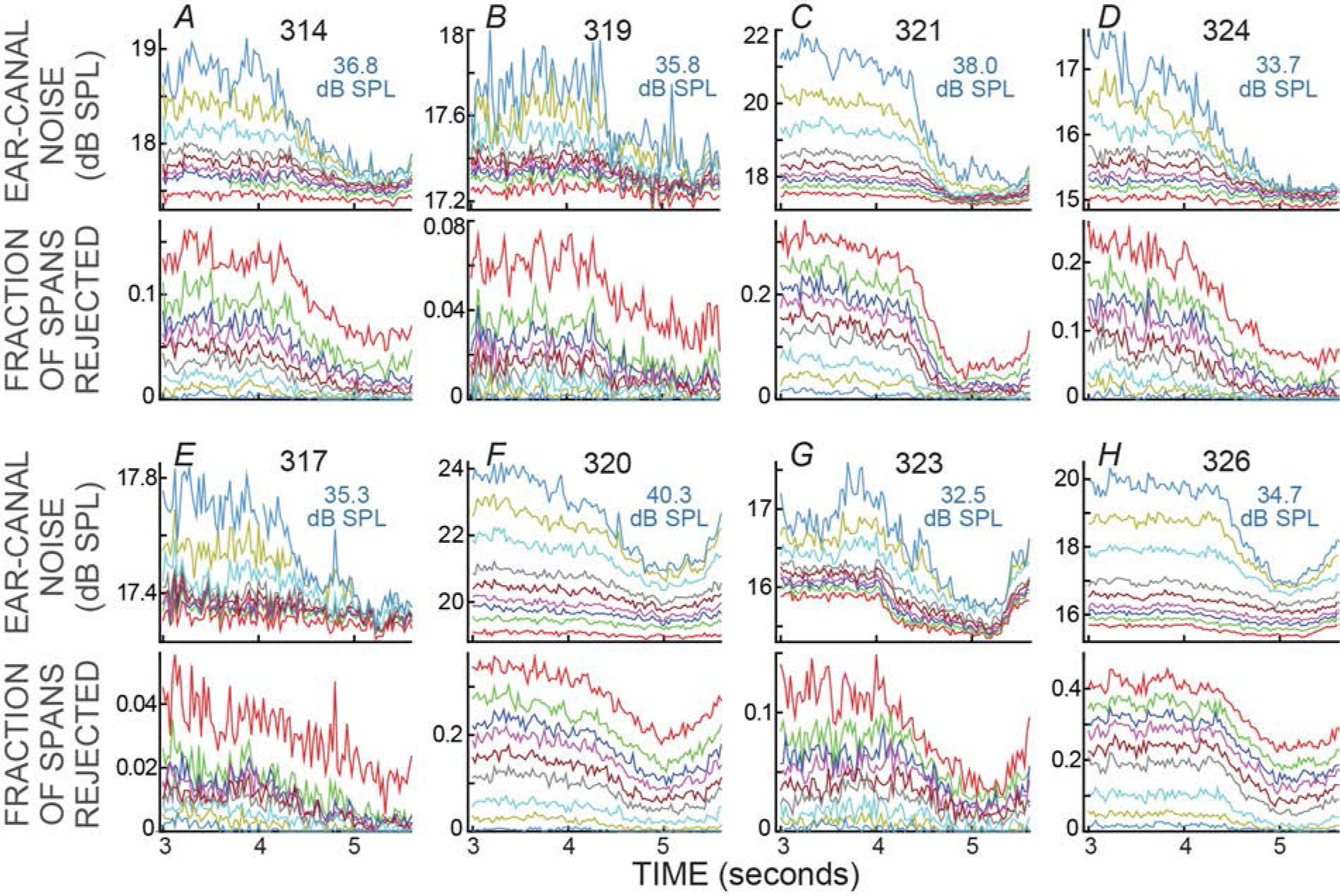
Ear-canal noise and fraction of spans rejected for the active trials of all subjects, for various edge-multiplier values. ***A-H***: For each subject there are two sub-panels (one above the other) with the subject number at top. The light blue text shows the sound-level cut-off value in dB SPL for an edge multiplier of 100. The edge multipliers were 2, 3, 4, 5, 7, 10, 20, 40, 100 (red to light blue lines, respectively; the sequence is the same in every panel and is most easily seen in the lower right panel). Note that as the edge multiplier decreased, more spans were rejected (the lines moved up in the lower panels) and the remaining ear-canal noise level was reduced (the lines moved down in the upper panels), but the shapes of the curves versus time remained similar.

The changes in ear-canal noise as the edge multiplier was changed from zero to 100, quantified as the change from epoch 10 to epoch 13, are shown in Figure 7. An edge-multiplier of zero applies a noise rejection criterion at the peak of the histogram of span RMS levels (see Fig. 2). Also included in Figure 7 are the changes from epoch 10 to epoch 13 of the raw data (data with no noise rejection applied). As the edge-multiplier was made less strict (i.e. had higher values) and fewer spans were rejected, the changes between epoch 10, and epoch 13 became larger for all subjects in active trials, but remained small in passive trials (Fig. 7A, B). To determine the extent to which the ear-canal noise was reduced more in active trials than passive trials, the difference between the two conditions is shown in Figure 7C. The difference was large when the edge multiplier was high and removed only the highest-level noise bursts, but as the edge multiplier was made more strict, the difference between the active and passive trials became less and less (Fig. 7C). For edge multipliers less than 1 there was almost no additional decrease (the decreases were less than 1%) in ear-canal noise produced in the active trials compared to the passive trials (Fig. 7C, inset). Note that using severe criteria (edge multipliers of 1 or less) did not remove the ability to see the small reductions in ear-canal noise in subjects 323 and 326 (Fig. 7A, B) - reductions that we attribute to the masking noise evoking MOC activity that reduced SOAEs and other noise of cochlear origin in these two subjects (Fig. 7B).

**Figure 7.**
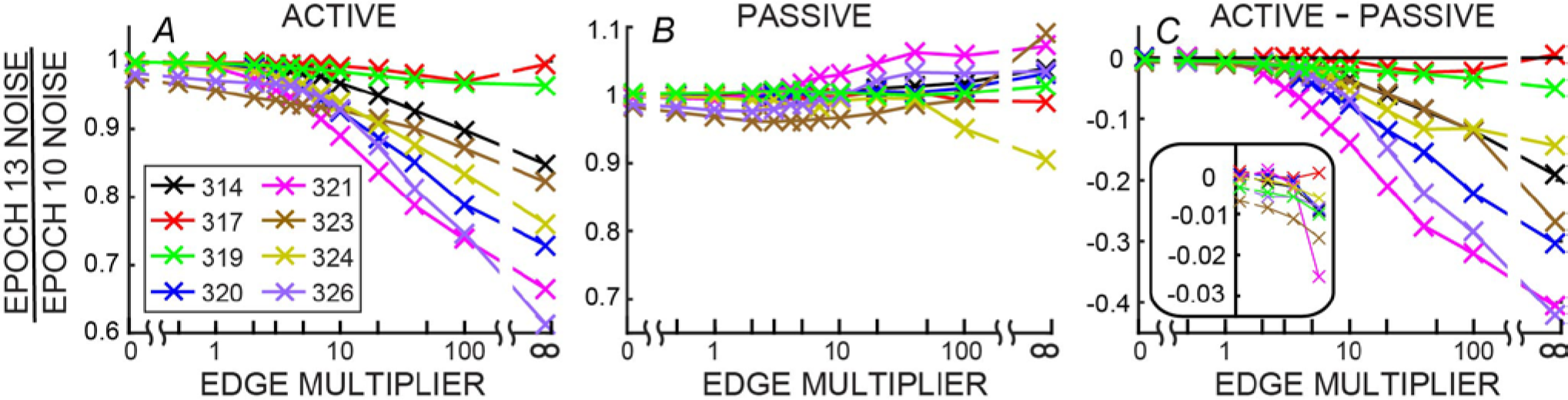
The change in ear-canal noise from epoch 10 to epoch 13 as a function of the edge multiplier for active trials (***A***), passive trials (***B***) and active trials minus passive trials(C) for each subject (key in box at left). The edge multiplier infinity sign indicates that no noise cut was done. The inset in C shows the lowest four points from each subject with an expanded vertical axis.

### 3.4 Noise spectra

Although the overall noise levels varied across subjects, all subjects showed similar patterns of ear-canal noise as a function of frequency. The noise amplitudes were largest at the lowest frequencies, were smallest at mid frequencies (2-3 kHz) and increased at higher frequencies (solid lines in Fig. 8A, B). The decrease from the original spectra to the spectra after applying an edge-extender of 2 was greater as frequency decreased (dashed lines in Fig. 8A, B). After noise bursts were removed by applying an edge multiplier of 2, there was little change in ear-canal noise from epoch 10 to epoch 13 at most frequencies (Fig. 8C, D). However, in the two subjects who showed reductions in SOAEs and/or other ear-canal noise with a time course appropriate for a MOC inhibition (subjects 323 and 326), there were decreases in the 1 to 2 kHz range (Fig. 8C, D). This frequency range approximately corresponds to the frequencies of these subjects’ SOAEs (Fig. 8E, F) and is also consistent with these changes being due to MOC inhibition.

**Figure 8.**
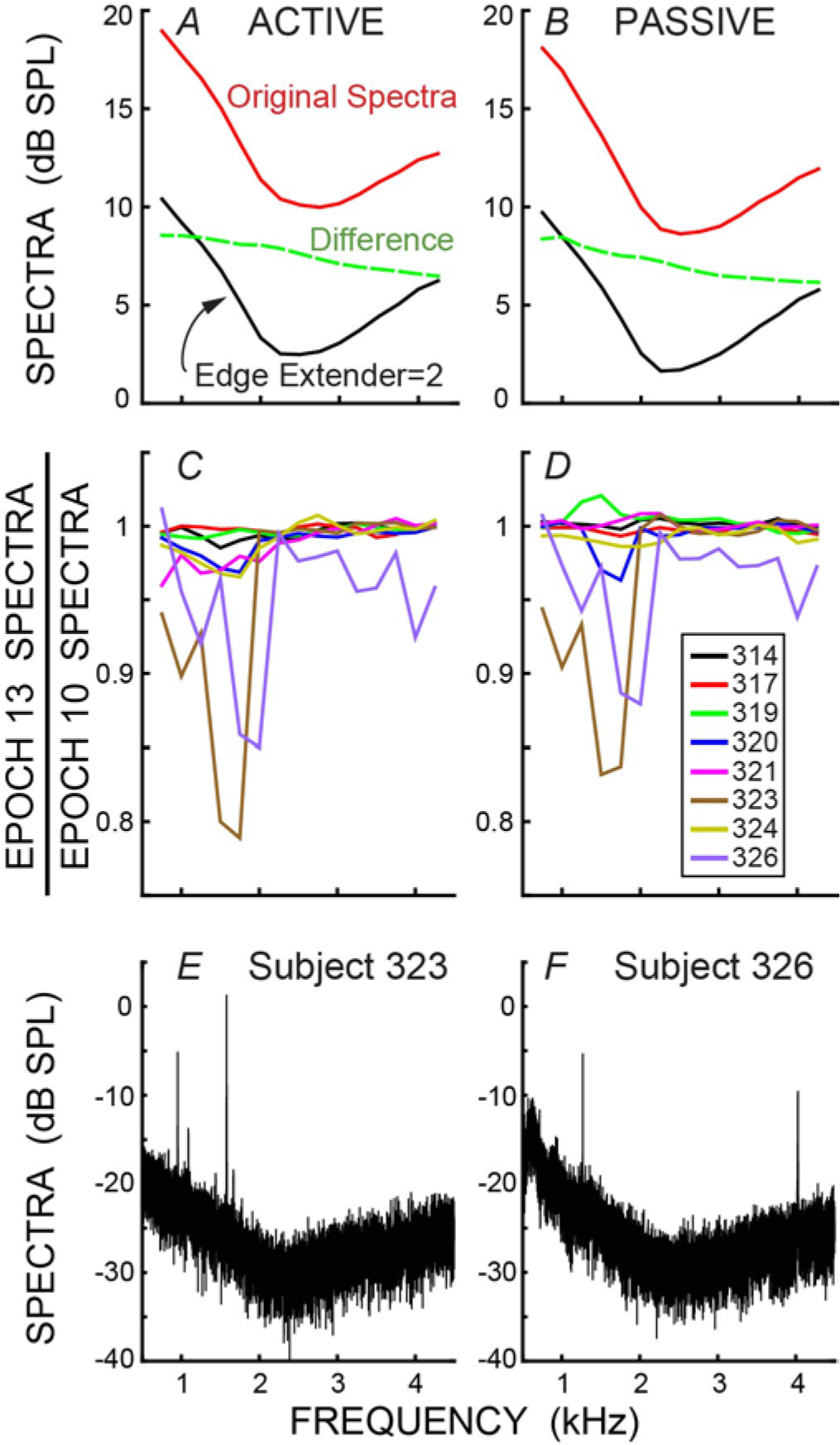
***A, B***: The spectra of the ear-canal noise, averaged across subjects, showing the original, un-cut spectra (top line) and the spectra after applying an edge-extender of 2 to remove spans with excess noise (bottom line). The dashed line is the difference (in dB, not SPL). ***C, D***: Data for an edge-multiplier of 2 showing the change in spectra from the baseline (epoch 10) to the last epoch during the masker noise (epoch 13) for individual subjects (key in box). *A, C* are for active trials; *B, D* are for passive trials. ***E, F***: Finely binned spectra showing the SOAEs in the two subjects who had SOAEs

## 4 Discussion

During the behavioral task we found reductions in ear-canal acoustic noise that were very large when no noise bursts were rejected, but became small when a strict criterion was used that removed most of the bursts of noise. The largest reductions in ear-canal noise were for active trials. We attribute the reductions in ear-canal noise as being due to two main sources: (1) inhibition from MOC efferent activity elicited by the task-ear masker noise, and (2) a reduction in subject motion concurrent with the subject attending to the task.

### 4.1 Reduction of ear-canal noise from MOC inhibition elicited by contralateral sound

A standard way of measuring MOC inhibition on OAE responses in one ear (here called the ipsilateral ear) has been to elicit MOC activity by contralateral acoustic stimulation (CAS). In the passive trials we did a measurement like that with the CAS being the task-ear masker. One difference from a typical MOC-effect measurement was that instead of measuring the effect on a sound-evoked OAE, we measured the effect on ear-canal acoustic “noise” (i.e. sound within the ear canal that was not evoked by a presented sound). In two subjects (323 and 326) we found strong evidence for reductions in ear-canal noise produced by CAS-elicited MOC inhibition: (1) the time courses of the reductions followed the typical time course of MOC inhibition produced by contralateral sound (Fig. 5), (2) as the criteria for removing ear-canal noise were made more strict, the changes from epoch 10 to epoch 13 did not go away, consistent with these changes *not* being due to changes in subject motion (Fig. 7A, B), and (3) the changes were found in both passive and active trials (Fig. 5C, D). These data fit with the hypothesis that in these two subjects, some of the ear-canal noise originated in the cochlea, and that MOC activity elicited by the masker CAS reduced cochlear amplifier gain thereby reducing the ear-canal noise. These two subjects were also the only subjects who had SOAEs, and it seems likely that much, or all, of the change was due to MOC inhibition of SOAEs (Mott et al., 1989; Harrison and Burns, 1993; Zhao and Dhar, 2010). However, it is also possible that some fraction of the change was actually MOC inhibition of a random signal that originated within the cochlea. Consistent with the hypothesis that some ear-canal noise in humans originates in the cochlea, Nuttall et al. (1997) found that basilar membrane velocity noise was enhanced by cochlear amplification and inhibited by MOC stimulation. This basilar membrane velocity noise can be expected to create backward-traveling noise waves that produce noise in the ear canal.

In addition to the two subjects with easily-seen decreases in ear-canal noise in passive trials, three other subjects also had very small, but statistically-significant decreases in ear-canal noise from epoch 10 to epoch 13. These may also have been MOC inhibitions of ear-canal noise or SOAEs that were too small to see. Overall, our finding of little or no CAS-elicited reduction in the ear-canal noise in subjects with no SOAEs is consistent with the hypothesis that in subjects with no SOAEs there is little or no noise in the ear canal that originated from within the cochlea.

The data without any noise rejection (Fig. 4) provide clear evidence that subjects reduced their ear-canal noise at the time the task was done. Several lines of evidence indicate that this was caused mostly by reduced subject motion, and not by task-elicited MOC activity reducing noise that originated within the cochlea. First, the largest noise bursts seem highly likely to have been produced by subject motion because their amplitudes are too high to be accounted for by any known cochlear mechanism. This is consistent with the normal interpretation in OAE measurements that large bursts of noise are due to subject motion. Second, when a strict criterion for removing large-amplitude noises was applied (e.g. an edge multiplier of 2 or less) there was almost no additional reduction in ear-canal noise in the active trials compared to the passive trials (Fig. 7C). Finally, one might think that attention-elicited MOC activity that reduced ear-canal noise would lead to a reduction of the number of spans rejected at that time. This explanation might then account for the pattern in Figure 6 where the reductions in the ear-canal noise and in the number of spans rejected have similar time courses. However, this explanation doesn’t fit with there being big reductions in ear-canal noise when the noise cut-off criterion was high (large edge multipliers), and small reductions as the cut-off criterion was made stricter. This pattern implies that when the subject did the task, the largest noise bursts were reduced more than the smallest noise bursts. For this pattern to be produced by MOC inhibition, the largest noise bursts would have to be inhibited more than the smaller noise bursts, which is opposite the pattern actually found for MOC inhibition at these sound levels (Guinan and Gifford, 1988; Guinan and Stankovic, 1996; Cooper and Guinan, 2006; Bhagat and Carter, 2010). Thus, the hypothesis that attention reduces ear-canal noise through MOC inhibition doesn’t fit the data for most subjects. A hypothesis that fits the data more broadly is that when attending to the task, the subjects sat more still and generated fewer bursts of noise.

It is interesting that the two subjects who showed clear evidence for CAS-elicited MOC inhibition of ear canal noise (323 & 326) also had slightly more change from epoch 10 to epoch 13 in active compared to passive trials (~1-2% greater during active trials; Fig. 7C, inset). One interpretation of this is that in these two subjects, task-related attention slightly increased the MOC activity and thereby produced a slightly greater epoch 10 to epoch 13 change in the active trials. However, since these changes were so small and absent in 6/8 subjects, we do not conclude that attention reduces ear-canal noise through MOC inhibition.

### 4.2 Comparison with previous reports

de Boer and Thornton (2007) reported reductions in ear-canal noise level when subjects did an auditory task or paid attention to a movie. They interpreted the changes they found in ear-canal noise as due to changes in subject-generated noise that were affected by attention and were also affected by whether the subject noise interfered with performance of the task. Their interpretation is consistent with ours.

In contrast, Walsh et al. (2014; 2015) reported a large decrease (~3 dB) in ear-canal noise in all of their subjects when the subject did a behavioral discrimination compared to during passive listening. They interpreted the decrease as being produced by MOC inhibition of ear-canal noise. The interpretation that this change was due to MOC inhibition is questionable for several reasons. A 3 dB reduction is at the high end of typical MOC effects on OAEs (Guinan, 2006) and would imply that a very large fraction of the ear-canal noise in *all* of their subjects originated within the cochlea, and also that there was a large attention-elicited MOC activation in the ear opposite to the task ear. Walsh et al. (2014; 2015) pointed out that a large attention-elicited MOC activation in the ear opposite to the task ear was unexpected because such efferent activation doesn’t help in performing the task. A second reason for questioning Walsh et al.’s interpretation is that their supposed MOC inhibition changed very little across frequency and was smallest in the 1-2 kHz range. Although there is some inconsistency across reports, previous work has always found that MOC effects are much greater in some frequency regions (often 1-2 kHz) than in others (Liberman, 1989; Veuillet et al. 1991; Chéry-Croze et al. 1993; Lilaonitkul and Guinan 2009a, 2009b, 2012; Zhao and Dhar, 2010, 2012). We think that the most economical hypothesis is that the large reductions of ear-canal acoustic noise reported by Walsh et al. (2014a; 2014b) were due to reductions in subject motion as the subjects attended to the tasks. However, there are many differences between Walsh et al.’s experiments and ours, so other factors cannot be ruled out.

### 4.3 Implications for measuring cochlear-efferent function with OAEs

Our results present a challenge for all experiments that seek, or have sought, to determine MOC activation by measuring OAEs during a behavioral task (Puel et al., 1988; Froehlich et al., 1990, 1993; Avan and Bonfils, 1988; Meric and Collet, 1992, 1993, 1994, 1996; Giard et al., 1994; Ferber-Viart et al., 1995; Maison et al., 2001; de Boer and Thornton, 2007; Harkrider and Bowers). It is typically assumed that when time periods containing large bursts of noise are removed, what remains is unaffected by subject motion. Our measurements indicate that no matter what level of artifact rejection was used, the ear-canal noise that remained was still affected by subject-generated ear canal noise. The simplest explanation is that rejection of large-amplitude “artifacts” does not remove all of the noise produced by subject motion or other physiological processes such as breathing. Although some ear-canal noise may originate from within the cochlea, an efferent effect on this noise is not easily separated from a similar-looking effect produced by decreased subject motion. This makes it difficult of measure MOC effects on ear-canal noise during a psychophysical experiment. In contrast, evoked OAEs, because they are similar from one trial to the next, can be separated from noise by averaging. Our results emphasize the need to have high signal-to-noise ratios (SNRs) in measurements of MOC-induced changes in OAEs (see Figure 8 of Goodman et al., 2013), particularly in behavioral experiments when subjects may decrease their movements and change ear-canal noise. The SNR needs to be high enough that changes in ear-canal noise will have a negligible effect on the signal measurement. In addition, our results also show that using a change in SNR as indicating there was a change in efferent inhibition (e.g. Sininger and Cone-Wesson, 2004) is not valid because the change could have been from changes in the noise. To sort out what part of ear-canal noise might be due to subject movement, it would be highly desirable to have an independent measure of subject movement, for instance a sensor attached to the head that tracks movements across time. However, simply showing the time course of the overall sound level (including noise and with no artifact rejection) during the experiment (as in Fig. 4) does provide a way of showing changes in subject-generated noise over time.

### 4.4 The reduction of ear-canal noise shows the time course of attention

In our behavioral 2IFC experiment, the onset of the masker noise was the only reliable timing cue that the tones were about to be presented, and that subjects should prepare to listen and give a behavioral response. Consistent with this cue timing, the reductions in the subjects’ ear-canal noise began approximately at the start of the masker noise (Figs. 4-6). In the absence of MOC inhibition, the time course of reductions in ear-canal noise, and in span rejections, can be thought of as indicators showing time course of subject’s directing their attention to the task. Although the time courses of the changes varied somewhat across subjects, the largest values were always near the time when the 2IFC target tones were presented (compare Figs. 1 and 6). The time course of the reduction in ear-canal noise shows that the buildup and decay of attention occurred over several hundreds of ms.

While we did not find that attention changed ear-canal noise through MOC inhibition, we did find that the decrease in ear-canal noise was a very robust indicator of whether subjects were or were not attending to sound. The time course of the decrease in ear-canal noise mirrors the time course of other physiological indicators of the preparatory control of attention. For example, Jaramillo and Zador (2011) found that during an auditory task, neural responses in auditory cortex increased as the expected moment of a target sound approached. Similarly, both pupil dilation (Irons et al, 2017) and neural activity in visual cortex (Stokes et al, 2009) show a rising time-course of activity that indicates the preparatory control of attention over a time-scale of seconds before executing a behavioral response to a visual target.

Recently, Gruters et al (2018) found an interaction between saccadic eye movement and changes in ear-canal sound pressure that lasted for 10’s of milliseconds. The infrasounds produced by such eye movements would have been filtered out in our measurements, but they do point out that there are many subject motion changes that may affect ear-canal noise. In addition, Braga et al (2016) found that saccade rates decrease during auditory attention. Thus, it is possible that as subjects attended to the auditory task, saccadic eye movements settled down, and this has a role in reducing ear-canal noise. If true, this hypothesis would indicate that eye-tracking might also help to sort out the origin of changes in ear-canal noise during task performance.

Our results indicate that before making definitive conclusions about the origin of changes in OAEs or ear-canal noise measured during a behavioral task, it is necessary to take into account all other sources that may affect ear-canal sound levels. This is especially true when studying MOC efferent effects, since extremely subtle motion artifacts may closely resemble MOC effects yet not be related to MOC inhibition.

## 6 Author Contributions

NF and JG: Designed the experiment; NF: Performed the experiment. NF, WZ and JG analyzed the data; NF and JG: Interpreted the results and wrote the manuscript. All authors read and approved the final manuscript.

